# Robust Brain State Decoding using Bidirectional Long Short Term Memory Networks in functional MRI

**DOI:** 10.1101/2021.06.18.449069

**Authors:** Anant Mittal, Priya Aggarwal, Luiz Pessoa, Anubha Gupta

## Abstract

Decoding brain states of the underlying cognitive processes via learning discriminative feature representations has recently gained a lot of interest in brain imaging studies. Particularly, there has been an impetus to encode the dynamics of brain functioning by analyzing temporal information available in the fMRI data. Long-short term memory (LSTM), a class of machine learning model possessing a “memory” component, is increasingly being observed to perform well in various applications with dynamic temporal behavior, including brain state decoding. Because of the dynamics and inherent latency in fMRI BOLD responses, future temporal context is crucial. However, it is neither encoded nor captured by the conventional LSTM model. This paper performs robust brain state decoding via information encapsulation from both the past and future instances of fMRI data via bi-directional LSTM. This allows for explicitly modeling the dynamics of BOLD response without any delay adjustment. The two hidden activations of forward and reverse directions in bi-LSTM are collated to build the “memory” of the model and are used to robustly predict the brain states at every time instance. Working memory data from the Human Connectome Project (HCP) is utilized for validation and was observed to perform 18% better than it’s unidirectional counterpart in terms of accuracy in predicting the brain states.

## 1 Introduction

Learning informative and discriminative representations of the brain states’ underlying various cognitive processes has gained a lot of interest in Brain-computer interface (BCI) applications [11]. Advances in non-invasive neuroimaging methods such as functional Magnetic Resonance Imaging (fMRI) are proving helpful in determining person’s cognitive or perceptual state [16]. e.g. in decoding motor functions [5], in the classification of shifts in attention [6], and for “brain reading” [3]. As a result, several techniques have been proposed for carrying-out brain-state decoding, from multi voxel-pattern analysis to understanding the behavior using deep learning architectures by integrating the spatio-temporal information.

Conventional decoding methods involved massive univariate analysis measuring activity from thousands of brain locations, analysing each of them separately [14]. The multivariate analysis takes into account the brain activity occurring at several locations simultaneously. This helps in integrating the distributed but overlapping information across the spatial domain [10]. Recent advances in time-sensitive machine learning frameworks have attracted remarkable attention for sequential modelling. In particular, two variations of the general Recurrent Neural Networks (RNNs) [20], namely, Echo-state Networks [19] and Long short-term memory models [12] have shown to perform better than conventional decoding models in characterizing dynamic fMRI information during both naturalistic and tasks conditions.

During the acquisition of fMRI data, the ratio of oxygenated to de-oxygenated blood level at any location in the brain serves as the representative of the underlying neuronal activation. Due to the time-lag observed in the peak of blood oxygen level dependent (BOLD) response, it is typically not considered to be synchronized with the presentation of stimuli [1]. Thus, in general, before training any brain-state decoding model, each time point is adjusted according to the the estimated delay of the BOLD signal [13], assuming that all fMRI voxels have the same response delay [17]. Long short-term memory (LSTM) [9], a class of RNNs, have been shown to model the temporal dynamic behaviour well. An LSTM model stores the information from past that has already passed through it and uses it as the contextual information for learning robust features for the intended task, say classification. Recently, some fMRI studies have used these networks for integrating the temporal information from past [12,19].

Because of the variations in latency of the BOLD responses across time, we assert that the temporal context from future is also important for capturing the dynamics of BOLD response in order to generate accurate representations. In this paper, we have employed a variant of LSTM architecture called Bidirectional LSTM [18], which acquires the information from both the past and future time-instances. In particular, the input sequence is fed in the normal time-order for one LSTM network and in the reverse time-order for another. The two hidden activations are collated to generate the hidden cell state features of the RNN. We evaluated this method for predicting brain states in working memory fMRI data obtained from the Human Connectome Project (HCP) [8]. The performance of the bi-LSTM network has been compared with its conventional unidirectional counterpart in brain state decoding task. This is to further note that this framework does not require any time-delay adjustment for the synchronization of stimuli and BOLD response unlike the previous works.

## 2 Materials and Method

### 2.1 Data

We evaluated the bi-LSTM framework on task fMRI data of the working memory from Human Connectome Project (HCP) [8]. We randomly selected *N* = 400 participants from the *N* = 1200 data release. Participants performed a working memory task, indicating if the current stimulus matches with the one presented two stimuli before, called “2-back” task, or a control condition called “0-back”. The working memory task from HCP also combines the category representation task. Hence, participants were presented with separate blocks of trials consisting of 4 different types of stimuli namely tools, places, faces, and body parts, separated with the fixation period. Data for two runs is available for each participant. Within each run, there were 8 task blocks for every task (2-back or 0-back) and stimuli (places, tools, faces, body) combination, each lasting for 27.5 seconds with 4 fixation blocks of 15 seconds each after two task blocks. Thus, each scan is a 405 time-points long sequence of fMRI volumes. More details about fMRI data acquisition and task paradigm are available at [8].

### 2.2 Data Preprocessing

The available preprocessed data [8] contains field-map based distortion correction, functional to structural alignment, and intensity normalization. Additionally, motion-related variables (6 translation parameters and their derivatives) were regressed-out using the *3dDeconvolve* with “ortvec” option in the AFNI software[4]. Changes in low frequency signals were regressed out using *3dDeconvolve* routine with the “polort” option. Since our goal is to evaluate a general brain-state decoding methodology, we used only the cortical data, which is directly available in surface representation as a part of HCP preprocessing pipeline. To separate brain areas based on architecture and functional connectivity, we employed the cortical parcellation developed by [7]. The parcellation method collates the individual voxels within each region by averaging to generate 360 cortical regions of interest (ROIs). The region-averaged timeseries was used as the input feature vector for the temporal analysis. The generated 405 time-points sequence of 360 ROIs were structured in a 360 by 405 2D-tensor. No stimuli and brain-state synchronization was performed to adjust for delay in the BOLD response in bi-LSTM network. Each time-point in task blocks was marked as present in one of the above mentioned brain-states and the time instants belonging to the fixation blocks were labelled as “others”, yielding a total of *S* = 9 brain states.

### 2.3 Bidirectional Long short-term memory RNNs

Brain-state decoding is essentially modelled as the task of classifying the brain state. Given the time-series of ROI brain features *x_t_* at time *t*, RNN model predicts the brain state of each time point based on input activation, *x_t_* and temporal dependency on its preceding time points until time *t* − 1. The LSTM, in particular, defines gated cells that can act on the received input activation by passing or blocking the information based on the importance of the feature for task. The learning process, called Back-propagation through time (BPTT) [20], estimates the parameters, which allow the data in the cells either to be retained or deleted. The transition equations for a LSTM model are as follows:

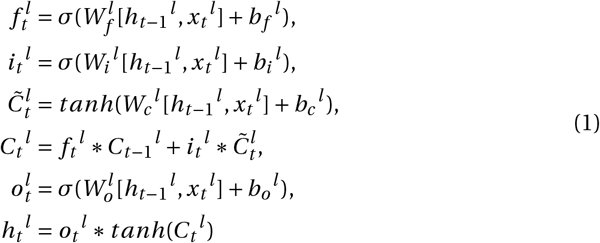

where *f_t_^l^*, *i_t_^l^*, *C_t_^l^*, and *x_t_^l^* denote the output of forget gate, input gate, cell activation, hidden activation, and the input activation of the *l^th^* LSTM layer at time point *t*. *σ* denotes the sigmoid activation function. A schematic illustration of Bidirectional LSTM [18] is provided in Fig. 1. It processes the time-series data in both the directions using separate hidden layers. It computes the forward hidden activation 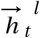 using the above equations. The backward hidden activation 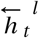 is calculated by using the future temporal dependency *h*_*t*+1_^*l*^. The input to the subsequent layer is generated by combining both forward and backward hidden activations as:

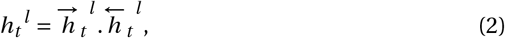

where dot (.) represents a merging scheme. The three possible ways of combining the activations are vector concatenation (bi-LSTM-c), element-wise vector addition (bi-LSTM-a), and element-wise averaging (bi-LSTM-*μ*). The input activations to the layer *l* = 1 at time *t*, *x_t_*, are extracted from the 360 brain ROIs and the input to the subsequent LSTM layers *l* = 2,3,., *n* are the hidden activations of the previous *l* − 1^*th*^ layer. The last layer of bi-LSTM is followed by a fully-connected layer having *S* neurons and softmax activation, and is used to learn a mapping from the learned feature representations to the brain states as:

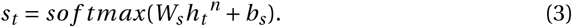

**Fig. 1.**
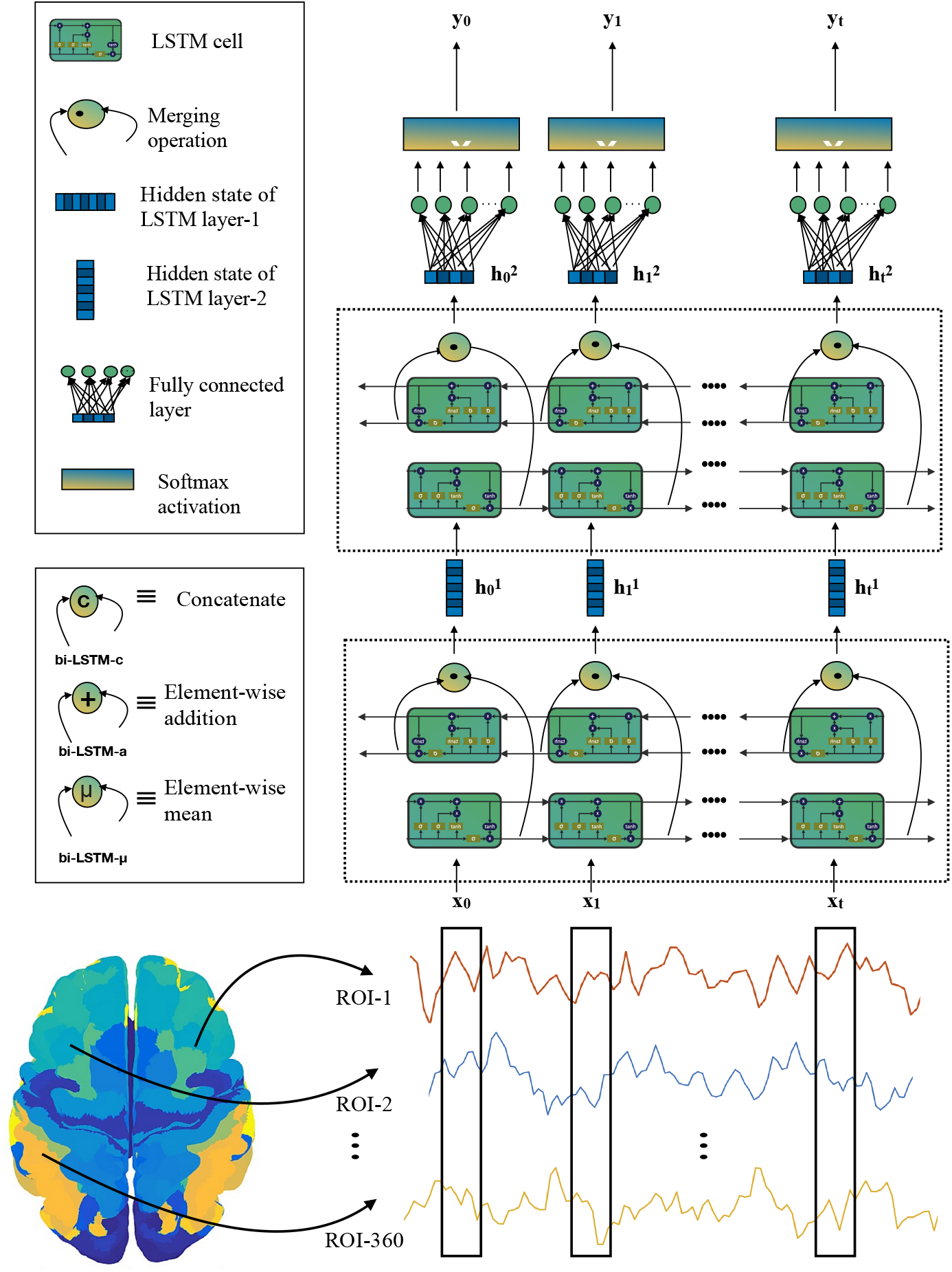
Schematic representation of the proposed framework for robust brain-state decoding using Bidirectional LSTM. At time *t*, 360 regions of interest (ROIs) are passed in as input activations, *x_t_*, to a 3-layer deep RNN architecture. The whole architecture has two stacked bidirectional layers for encoding temporal dependencies, followed by a fully-connected layer with softmax activation for predicting the brain-states. One bidirectional layer comprises of a set of forward and backward layer of LSTM, and is highlighted using a dashed-line box. The hidden activations from forward and backward LSTMs can be merged in three different ways. The merging schemes are depicted in a box in the middle left. Based on this, the models can be named as bi-LSTM-c, bi-LSTM-*μ*, and bi-LSTM-a. The output vector from the fully-connected layer *y_t_* indicates the predicted brain-state corresponding to the input *x_t_*.

## 3 Experimental Results and Discussion

For comparing the performance of the bi-LSTM with the conventional LSTM that was earlier evaluated on the working memory task, bi-LSTM architecture with the same specifications as in [12] was built. At any given time *t*, input activations *x_t_* pass through two hidden bi-LSTM layers with each LSTM cell having 256 hidden activations to encode the temporal dependencies. This is followed by a fully-connected layer containing *S* = 9 neurons predicting the brain states. We employed *inter-subject* 10-fold cross-validation. Data from *N* = 400 participants was divided into 10 parts, of which 9 parts (*N* = 360 subjects) were kept for model development and the remaining one part (*N* = 40 subjects) was kept unseen for evaluating the model performance. The development set (9 folds) was also randomly shuffled and split into 80-20:training-validation sets. The validation data was generated to tune the hyperparameters and to prevent over-fitting. Since the task-paradigm for each run and each subject was same during the acquisition of the working memory task data [12], the full-length training data was windowed into small overlapping sizes with the window size *w* = 40 with an overlap of 10 points [12].

The proposed model was implemented in Keras [2] deep learning framework. The model was trained on GeForce GTX 980 GPU with a batch size of 32, using ADAM optimizer with a learning rate of 0.001. The model was trained for 100 epochs and no early stopping was performed. To prevent the model from over-fitting, a dropout of 0.3 was applied in LSTM layers for training. The number of time-instances of class “others” in the data were much more than any other class. Thus, in order to prevent the model from predicting the states as per the underlying class distribution, weights for the imbalanced classes were estimated using Sklearn’s “compute_class_weight” [15] routine and were applied during loss function calculation, giving value to instances that was inversely proportional to their frequency in the data.

We compared the proposed architecture (bi-LSTM) with its conventional unidirectional counterpart LSTM and with a three layer feed-forward Neural Network (ff-NN), which used ROIs at individual time points as features discarding temporal dependencies. For better comparison, the number of layers and the number of neural units in the layers for the other models were kept same as in the proposed model. We also used different models of bi-LSTM Fig. 1 based on the combination of activations of the forward 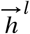 and backward 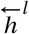 hidden states, namely, bi-LSTM performing merging by concatenation (bi-LSTM-c), element-wise adding (bi-LSTM-a), and by taking element-wise mean (bi-LSTM-*μ*). Results in Table-1 were obtained by evaluating the performance on the unseen test data of *N* = 40 subjects in each fold. The averaged *F*_1_ score for each class are tabulated in Table-1. It is observed that the bidirectional models outperform ff-NN and LSTM. Further, bi-LSTM-c seems to perform slightly better than the bi-LSTM-*μ* and bi-LSTM-a. Possibly, summation or averaging may be merging the features activations leading to slightly inferior performance.

**Table 1.** Comparative performance of different models in terms of cross-validated *F*_1_ score for each brain state and the weighted-average performance using the unseen data of 40 participants from working memory task fMRI data.

| Model | 0-back<br>Body | 0-back<br>Faces | 0-back<br>Places | 0-back<br>Tools | 2-back<br>Body | 2-back<br>Faces | 2-back<br>Places | 2-back<br>Tools | Others | Weighted<br>Average |
| --- | --- | --- | --- | --- | --- | --- | --- | --- | --- | --- |
| ff-NN | 0.53 | 0.54 | 0.52 | 0.48 | 0.48 | 0.60 | 0.53 | 0.52 | 0.79 | 0.55 |
| u-LSTM | 0.68 | 0.64 | 0.69 | 0.62 | 0.56 | 0.70 | 0.69 | 0.61 | 0.71 | 0.66 |
| bi-LSTM-μ | **0.85** | **0.83** | 0.86 | 0.81 | 0.79 | **0.87** | 0.86 | 0.84 | 0.87 | 0.85 |
| bi-LSTM-c | **0.85** | **0.83** | **0.87** | **0.83** | **0.80** | **0.87** | **0.87** | **0.85** | **0.88** | **0.86** |
| bi-LSTM-a | **0.85** | **0.83** | 0.85 | 0.81 | **0.80** | 0.86 | **0.87** | 0.85 | **0.87** | **0.85** |

Note: Best classification performance for each brain-state is highlighted in bold.

Mean normalised confusion matrices on the classification accuracy are illustrated in Fig. 2 for comparing the miss-classifications of both LSTM and bi-LSTM-c. The overall accuracy of the unidirectional LSTM model was 0.66 ±0.18, whereas the classification accuracy of the bidirectional LSTM (bi-LSTM-c) was 0.84 ±0.02. For every brain-state, the LSTM model miss-classify to a larger extent compared to the bi-LSTM-c model, although the misclassification is highest for both the models to the “Others” class. Furthermore, the second highest confusion in case when participants were stimulated with “faces” and “places” is with the task (0-back or 2-back), though the stimuli was detected correctly. The model also gets confused between the stimuli “body” and “tools”.

**Fig. 2.**
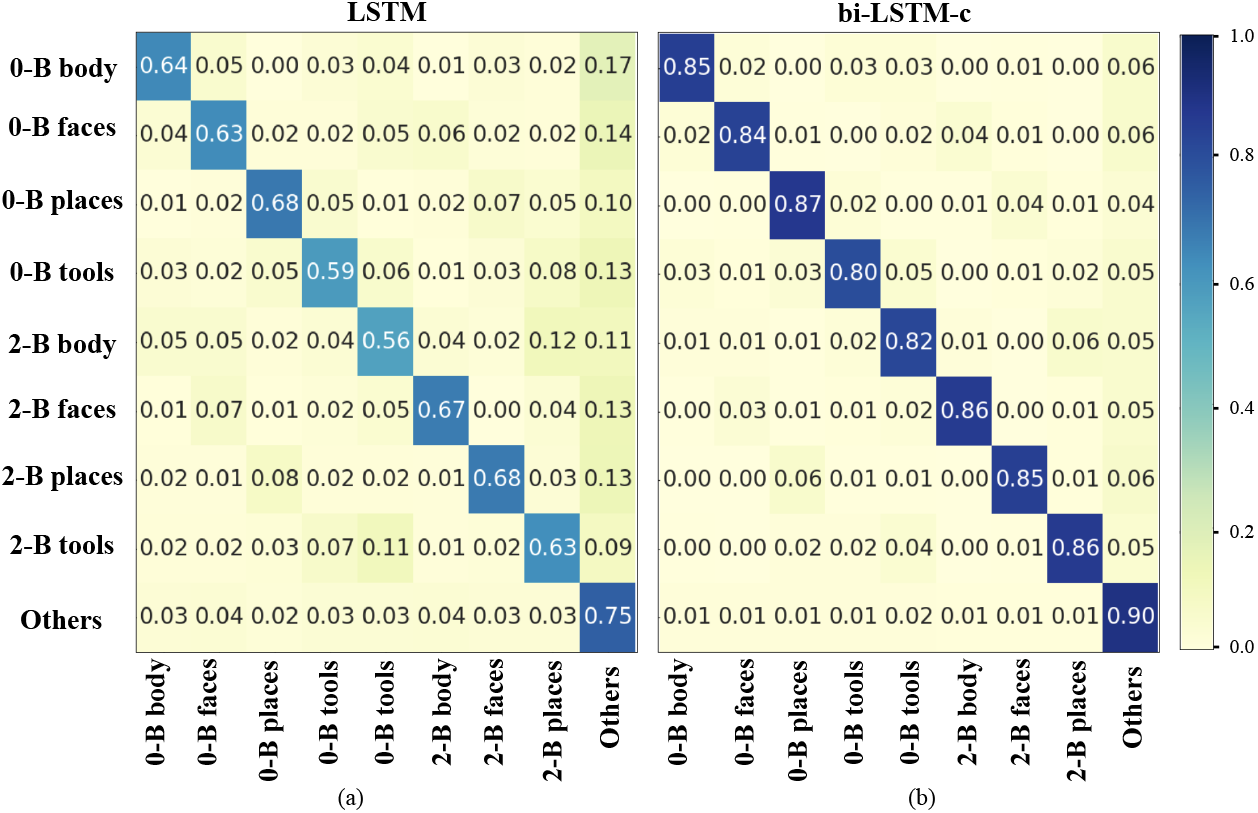
Brain-state decoding performance of (a) Long short-term memory (LSTM) and (b) Bidirectional LSTM with feature concatenated merging (bi-LSTM-c) on the unseen data of 40 particpants from working memory task fMRI data. The color bar indicates mean accuracy across 10 cross-folds of validation.

## 4 Conclusions and Future Work

In this study, we propose to use Bi-directional LSTM network model for decoding brain states from task fMRI data in order to appropriately capture the dynamics of fMRI BOLD response. The experimental results on the working memory task fMRI data demonstrated superior performance of Bi-LSTM compared to the unidirectional LSTM. Further, this model works well without any hard-coded delay adjustment, emphasizing the availability of useful information in the immediate future samples as well. We worked with the fixed window length, although future work may involve tuning the window-size and overlap for time-series chunking. The problem of class imbalance, although, majorly handled, still requires more sophisticated handling. From analysis point of view, it would be interesting to study about cortical regions engaged in stimuli “body” and “tools” as the model sometimes gets confused between them.

## Notes

### Competing Interest Statement

The authors have declared no competing interest.

### Summary of Updates

The manuscript is revised to correct the spelling mistake of one of the author. The author Luiz Pessoa's name in PDF version of manuscript was written as Luis Pessoa

